# Human Papillomavirus Type 16 E6 induces cell competition

**DOI:** 10.1101/2021.07.06.451240

**Authors:** Nicole Brimer, Scott Vande Pol

**Affiliations:** Department of Pathology, University of Virginia, Charlottesville, Virginia, USA

**Keywords:** HPV-16, p53, apoptosis, transformation, cancer, tumorigenicity

## Abstract

High risk human papillomavirus (HPV) infections induce squamous epithelial tumors in which the virus replicates. Initially, the virus-infected epithelial cells are untransformed, but expand in both number and area at the expense of normal squamous epithelial cells. How this occurs is unknown, but is presumed to be due to viral oncogene expression. We have developed an *in vitro* assay in which colonies of post-confluent HPV16 expressing cells outcompete confluent surrounding normal keratinocytes for surface area. The enhanced cell competition induced by the complete HPV16 genome is conferred by E6 expression alone, and not by individual expression of E5 or E7. In traditional oncogene assays, E7 is a more potent oncogene than E6, but such assays do not include interaction with normal surrounding cells. These new results separate classic oncogenicity that is primarily conferred by E7, from cell competition that we show is primarily conferred by E6, and provides a new biological role for E6 oncoproteins from high risk human papillomaviruses.

**Importance:** High risk papillomavirus infections induce epithelial tumors, some of which evolve into malignancies. The development and maintenance of cancer is due to the virally encoded E6 and E7 oncoproteins. How a virally infected keratinocyte out-competes normal uninfected keratinocytes has been unknown. The present work shows that the enhanced competition of HPV16-infected cells is primarily due to the expression of the E6 oncoprotein and not the E7 or E5 oncoproteins. This work shows the importance of measuring oncoprotein traits in the context of cell competition with uninfected cells, and shows the potential of papillomavirus oncoproteins to be novel genetic probes for the analysis of cell competition.

## Introduction

Papillomaviruses induce epithelial hyperplasias (papillomas) in vertebrates, that can vary in size from visually inapparent up to kilogram masses (1). The virus replicates in the papilloma under the control of virus-encoded E1 and E2 proteins (2, 3). The virally-encoded E5, E6, and E7 oncoproteins contribute to the formation of the papilloma (4–6), and are expressed under the transcriptional control of cellular transcription factors together with the E1 and E2 proteins (7–14). In some HPV types and in Bovine Papillomavirus type I, the complete papillomavirus replication cycle can be studied in vitro using keratinocyte organotypic culture and cloned viral DNA (15–19).

The papillomavirus infection cycle begins with exposure of basal epithelial cells and the basement membrane to a virus inoculum; virus associates with the basement membrane, is taken up by basal epithelial cells, and early genes including the viral oncoproteins are expressed (20). The initially infected cell(s) must attach to and persist on the basement membrane, because if the attachment is lost, initially virus infected cell(s) could be forced apically by other basal proliferating cells, resulting in the loss of the infected cell by desquamation from the epithelial surface. Therefore, attachment to the basement membrane and the ability of daughter cells to remain attached and proliferate at the expense of surrounding uninfected epithelium is a requirement for an incipient papilloma to expand. How virally infected cells compete at the expense of uninfected keratinocytes is presumably (but as yet unproven to be) a consequence of viral oncoprotein expression; but which oncoprotein(s) most influence cell competition is as yet unknown.

Cell competition is a rapidly expanding field that originated from the observation that normal cells in *Drosophila* embryos eliminate adjacent cells that are impaired but viable (due to having only a single copy of a ribosomal protein gene) ((21) and references therein). In cell competition, normal cells eliminate cells with reduced fitness. Described mechanisms include the induction of apoptosis, mechanical competition, and intercellular signaling that induces differentiation (recently reviewed in (22, 23)). While the hallmark of cell competition is normal cells eliminating impaired cells, abnormal super-competing cells can eliminate normal cells as is seen when super-competing cells produced by overexpression of c-myc can induce the displacement or death of normal surrounding cells (24, 25). Since papillomaviruses induce papillomas that expand at the expense of normal tissues, viral manipulation of cell competition may play a role. The viral oncoproteins E5, E6, and E7 would be candidates to manipulate cell competition. These viral oncoproteins have been characterized by classic oncoprotein assays such as focus formation, inducing anchorage independent growth of 3T3 cells, or transgenic expression in murine skin, all of which are all assays that measure the oncoprotein’s traits in homogenous cell populations. Such assays do not recapitulate the early stages of an in vivo infection where a virally infected cell population expands by successfully competing against normal cells for space to form a lesion.

In classic assays for oncogene activity, the major oncogene of HPV16 is 16E7, which when compared to 16E6, has increased ability to induce anchorage independent colonies (26, 27) and induces a more severe dysplasia than 16E6 when expressed in the skin of mice (28–30). The E7 oncoproteins are best known for targeting the degradation of Retinoblastoma family proteins (31–33) and the tyrosine phosphatase PTPN14 that is a negative regulator of Hippo signaling (34–36). If the ability of HPV16 genomes to induce enhanced cell competition were due to oncogenic potency, E7 would seem to be the most likely candidate. However, in experiments presented here we find that it is HPV16 E6 and not E7 or E5 that independently enhances cell competition.

## Materials and Methods

### Cell culture

NIKS Keratinocytes are human foreskin keratinocytes that are both feeder-cell and growth-factor dependent for proliferation, support the complete HPV lifecycle, are untransformed, and have an extended lifespan (37); they were obtained from ATCC (https://www.atcc.org). NIKS were co-cultured with mitomycin C treated 3T3 cells in F-media and transduced with replication defective lentiviruses and with replication defective murine retroviruses as previously described (38). NIKS cells were transfected with re-circularized cloned HPV16; episomal status of the HPV16 genome was confirmed by southern blot (39). Primary keratinocytes were derived from anonymous discarded neonatal foreskins collected from the University of Virginia Medical Center and classified as non-human subject research, and were maintained and virally transduced in F-media with mitomycin C treated 3T3 cells and Rho Kinase inhibitor (Y-27632, ThermoFisher) as previously described (40).

### Plasmids

HPV16 nts 56-879 encompassing the E6 and E7 region cloned into murine retrovirus vector pLXSN was the kind gift of Denise Galloway (41). Stop codons were introduced at amino acid 12 of E6 and amino acid 8 of E7 either alone or in combination as shown in the figures. HPV16 E5 cloned into a retroviral expression vector was the kind gift of Richard Schlegel (Georgetown University) (42). EGFP (from Clontech) or Fusion Red ((43) obtained from Addgene clone 54778) were cloned into a lentiviral packaging plasmid with an internal MSCV promoter and puromycin selection.

### Cell Competition Assay

Primary Keratinocytes cultured in F media in the presence of Y-27632, or NIKS cells, or HPV16 transfected NIKS cells, were cultured in F media with feeder cells as described above. Keratinocytes transduced with either the EGFP or Fusion Red lentivirus were then transduced with the above-described murine retroviruses expressing either wild-type or mutated 16E6, and/or 16E7 or 16E5 and drug-selected in F media with puromycin and G418 (and in the case of primary cells, 10 uM Y-27632). One day before the beginning of the assay, 99.5% to 99.9% vector-expressing cells and 0.1 to 0.5% oncogene-expressing cells in contrasting fluorescent tagged cells were mixed and plated together at 10% confluency in a 10 cm dish. 24 hours later, those cells were trypsinized and re-plated onto glass coverslips in a 6 well plate (at 2.1×10^4^ cells / cm2) together with mitomycin-C treated feeder 3T3 cells in F media (with or without Y27632). Cells typically reached confluency at day 5-7, at which point one well is fixed and a second well is fed on alternate days with F-media for another 7 days until fixation and a third well fixed on days 17-21. Coverslips were stained with dapi. Fluorescent images were acquired as 16-bit TIFF images with a Nikon inverted TE-2000-E fluorescence microscope equipped with a Retiga6 camera (Photometrics.com) controlled by Oculus software. Pictures of fluorescent colonies were taken from randomly selected fields and the relative size of the colonies ascertained using Fiji image analysis software (https://ImageJ.github.io).

### Western Blotting

SDS-lysed keratinocyte cell lysates were equalized for protein concentration (BioRad). Equal amounts of protein-normalized samples were loaded onto SDS-acrylamide gels, electrophoresed, and transferred onto PVDF membranes. Blots were blocked in 0.05% tween-20/5% Non-fat milk in Tris-buffered saline and probed with the indicated antibodies from Cell Signaling: GAPDH (#3683), Tubulin (#T9026); from BD Biosciences: anti-AU1 tag monoclonal antibody was a gift of Richard Schlegel (Georgetown University), Actin (ACTN05) (MS-1295-P1), p53 (MA1-19055); anti-16E6 mAb 6G6 was a kind gift of Johannes Schweitzer (Arbor Vita Corp.) and anti-16E7 was a mix of both monoclonal antibody clones 8C9 and EDV7 (Santa Cruz Biotechnology).

## Results

### HPV16 confers enhanced cell competition to keratinocytes

An assay was developed to measure the relative fitness of keratinocytes harboring HPV16 in competition with uninfected keratinocytes. The assay mimics the early stage of an HPV16 infection where an infected keratinocyte establishes a nascent papilloma and is illustrated in Fig. 1. 0.1 to 0.5% HPV16 transfected and fluorescently-tagged keratinocytes are seeded together with 99.5-99.9% vector-transduced keratinocytes fluorescently tagged with a contrasting colored protein at low density into a 10 cm dish for 24 hrs. before trypsinization and re-seeding of the mixed cell population together onto coverslips in a 6-well plate. The 24 hr. co-culture in the 10 cm plate insures that both populations begin the cell competition assay on glass coverslips under identical culture conditions. Confluency is reached about 5-7 days after seeding onto coverslips, at which time one coverslip is fixed and dapi stained, and the remaining coverslips are cultured for a further 7-14 days with feeding on alternate days (becoming super-confluent), and then are fixed and stained with dapi. Daily feeding of the coverslips leads to more rapid super-confluency and a shorter assay duration. A variety of fluorescent proteins were screened for this assay (EGFP, Fusion Red, mCherry, mCitrine, mVenus, and mCerulean) with EGFP and Fusion Red being chosen both for similar toxicity and spectral properties. The assay duration is limited by eventual stratification of epithelial cells into multicellular ridges that develop auto-fluorescence and interfere with fluorescence imaging. Pictures were taken of random fields and the relative size of colonies calculated. As super confluency is reached, the 3T3 feeder cells in the culture are forced off the plate by the keratinocytes and auto-fluoresce; these balls of auto-fluorescent cells were excluded from analysis.

**Fig.1.**
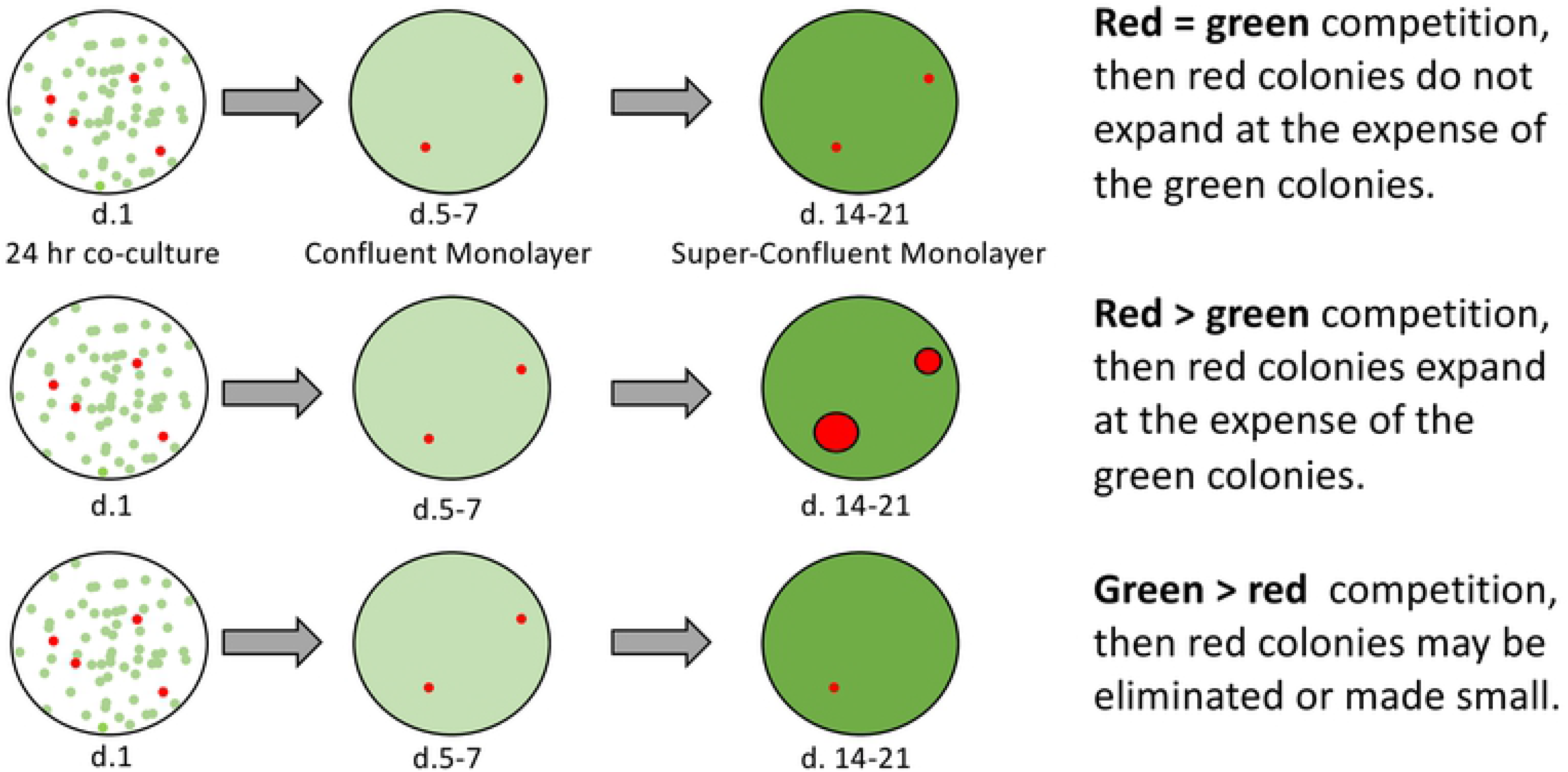
An in vitro model for papillomavirus infected cell competition. Keratinocytes were labeled by lentiviral transduction with either green (eGFP) or red (Fusion Red) proteins and then transduced with either vector or papillomavirus expressing retroviruses and cultured separately. On day 1 of the assay, 99.9% vector-expressing green cells and 0.1% oncogene expressing red cells were mixed and plated together at 10% confluency in a 10 cm dish. 24 hours later, those cells were trypsinized and each sample was plated onto 3 glass coverslips in a 6 well plate (2.1 10^4^ cells / cm^2^). Cells reach confluency by day 5-7 at which point one well is fixed and a second well is fed daily with F-media for another 7 days and then fixed, and a third well incubated for another 7 days before fixation, and staining with dapi. Pictures of fluorescent colonies were taken with a 4X objective from randomly selected fields and the relative size of the colonies ascertained using NIH ImageJ software.

If HPV16 conferred no competitive advantage, the colony sizes should not exceed those of vector transduced cells (Fig. 1), but that was not the case (Fig. 2). HPV16-expressing red keratinocyte colonies expanded in surface area at the expense of surrounding vector-transduced green cells, while red vector-transduced cells only modestly out-competed green cells (Fig. 2c, d, k).

**Fig.2.**
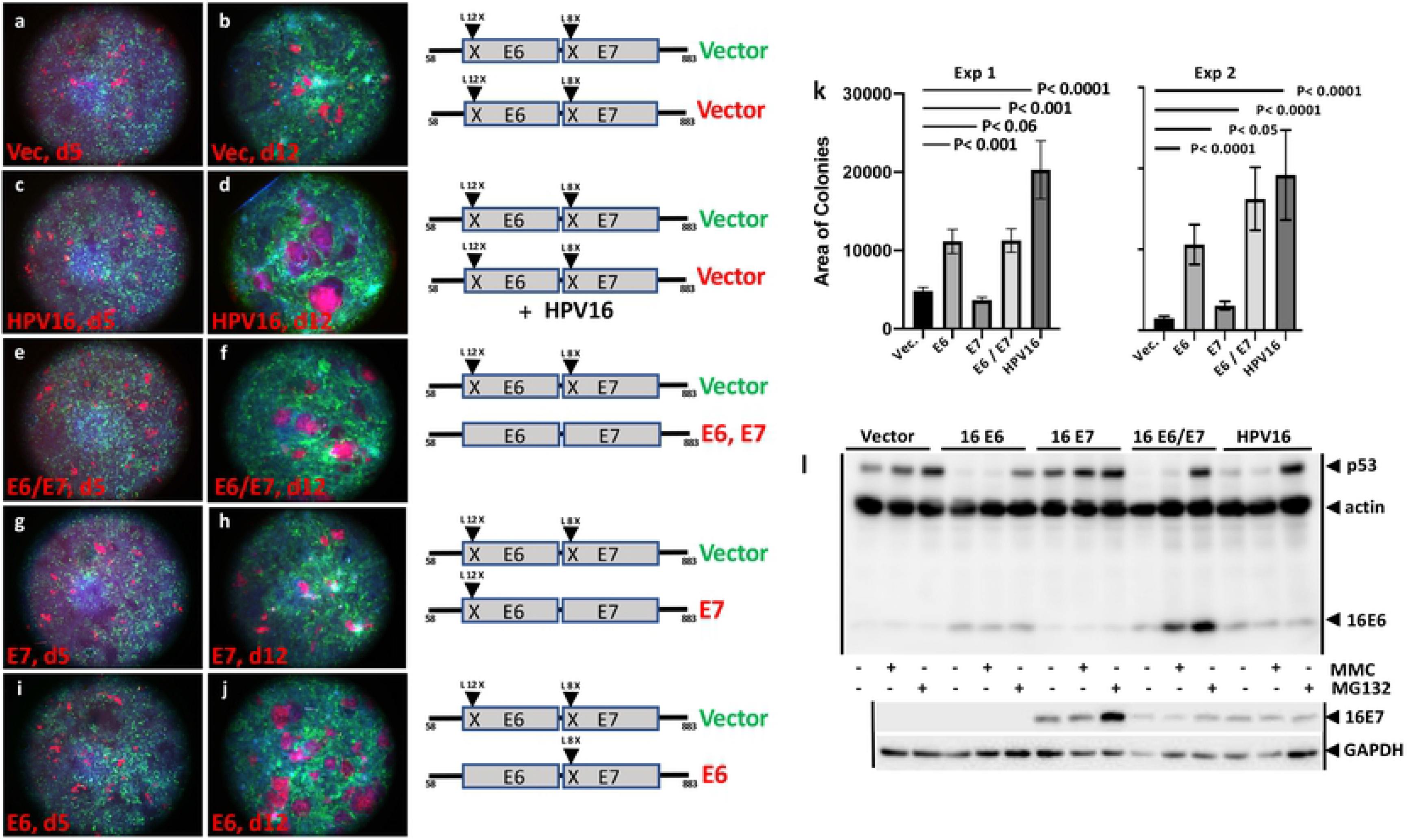
Enhanced cell competition is induced by HPV16 and by HPV16 E6. Vector-transduced green cells and oncogene-expressing red cells were seeded together as described in Fig. 1 onto coverslips on day 2 and fixed at confluency on day 5 (a, c, e, g, i); a second coverslip was fixed on day 12 (b, d, f, h, j). The transduced genes are indicated to the right of each pair. The quantified results of colony sizes (in arbitrary units) from 2 experiments is shown (k). Western blots for the stably transduced cell lines selected in Fusion Red expressing cells is shown (l). HPV16 and E6 confer enhanced cell colony size at day 12 while E7 does not. Day 5 colony sizes were not statistically different between the samples and are not shown. Error bars in part k depict standard error of the mean for colony sizes. The two assays shown are representative of 4 assays.

### E6 and E7 proteins phenocopy enhanced competition caused by the complete HPV16 genome

HPV16 encodes E5, E6, and E7 oncoproteins as well as RNA products that encode additional proteins and could have additional non-proteinaceous functions. In most cervical cancers, only the E6 and E7 genes are expressed after viral integration in the E2 or E1 genes (44, 45). In order to determine if only the E6 and E7 proteins are sufficient to confer enhanced cell competition, retroviral vectors expressing the HPV16 E6 and E7 proteins were introduced into red keratinocytes while an identical E6 and E7 expression vector with stop codons introduced early in the E6 and E7 ORFs was introduced into the green cells in order to insure that the red and green cells express common RNA products and drug selection markers, and differ only in the expression of E6 and E7 proteins and fluorescent markers. Fig. 2e, f, and k show that E6 together with E7 proteins alone conferred enhanced cell competition. Thus, neither E5 nor other virally encoded functions in HPV16 are essential for enhanced cell competition conferred by the HPV16 genome.

### 16E6 predominantly confers enhanced cell competition compared to E7

The indicated retroviral constructions in Fig. 2 were used to express only E6 or only E7 in red cells compared to green cells. While 16E6-expressing red keratinocytes outcompeted green vector-transduced keratinocytes, 16E7-expressing keratinocyte colonies were smaller than vector-transduced keratinocytes (although the smaller colony size did not reach statistical significance). The absence of enhanced cell competition by 16E7 was not due to an absence of E7 expression as both oncoproteins were expressed where expected and not where mutated (Fig. 2l). The differences in cell competition were quantified as differences in the size of the keratinocyte colonies as ascertained by automated quantification of randomly selected microscopic fields and reached high statistical significance (Fig. 2k).

To confirm that the results of Fig 2 were not the result of different fluorescent protein tags, the Fig. 2 assay was repeated with the oncoproteins expressed in green cells and the surrounding competing cells tagged with Fusion Red. Fig. 3 shows similar competition results regardless of green or red fluorescent tags employed.

**Fig. 3.**
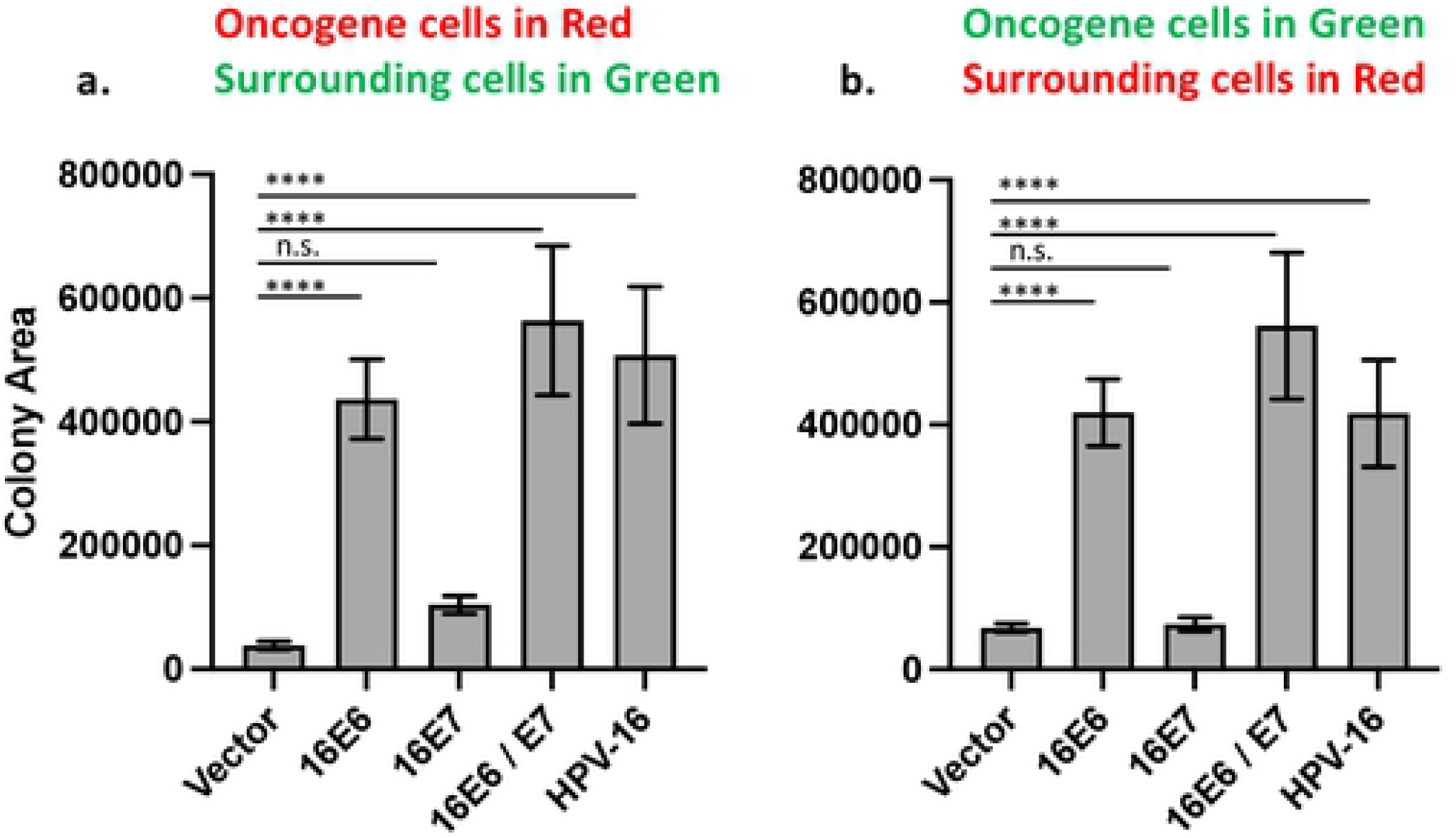
Cell competition induced by HPV16 or HPV16 E6 alone is similar in competing cells expressing either EGFP or Fusion Red tags. The same oncogene transductions shown in Fig. 2 were expressed in either Fusion Red (a) or EGFP tagged cells (b) and put into competition with the alternate colored cells as shown and described in Fig. 2. Colony sizes are shown in arbitrary units and error is standard error of the mean. **** is P<0.0001; n.s. is not significant.

### 16E6 alone confers enhanced competition

The ability of the 16E6 ORF alone to enhance cell competition did not distinguish between the full length E6 protein or a smaller E6 protein termed E6*, where the E6* mRNA encodes the first 41 amino acids of 16E6, splices and terminates two amino acids later. Additionally, the absence of the E5 ORF in the retroviral E6/E7 retrovirus used in Fig. 2 left out the possibility that E5 might enhance cell competition. To address this and confirm the validity of the Fig. 2 results, red keratinocytes were transduced with retroviruses expressing only the individual E5, E6*, E6, E7 and E6 with a stop codon at aa 12 (as the negative control) and set into competition with green keratinocytes transduced with the corresponding null-expression retroviral vector. Fig. 4 shows that 16E6 markedly increased cell competition while E6*, 16E5, and 16E7 did not.

**Fig. 4.**
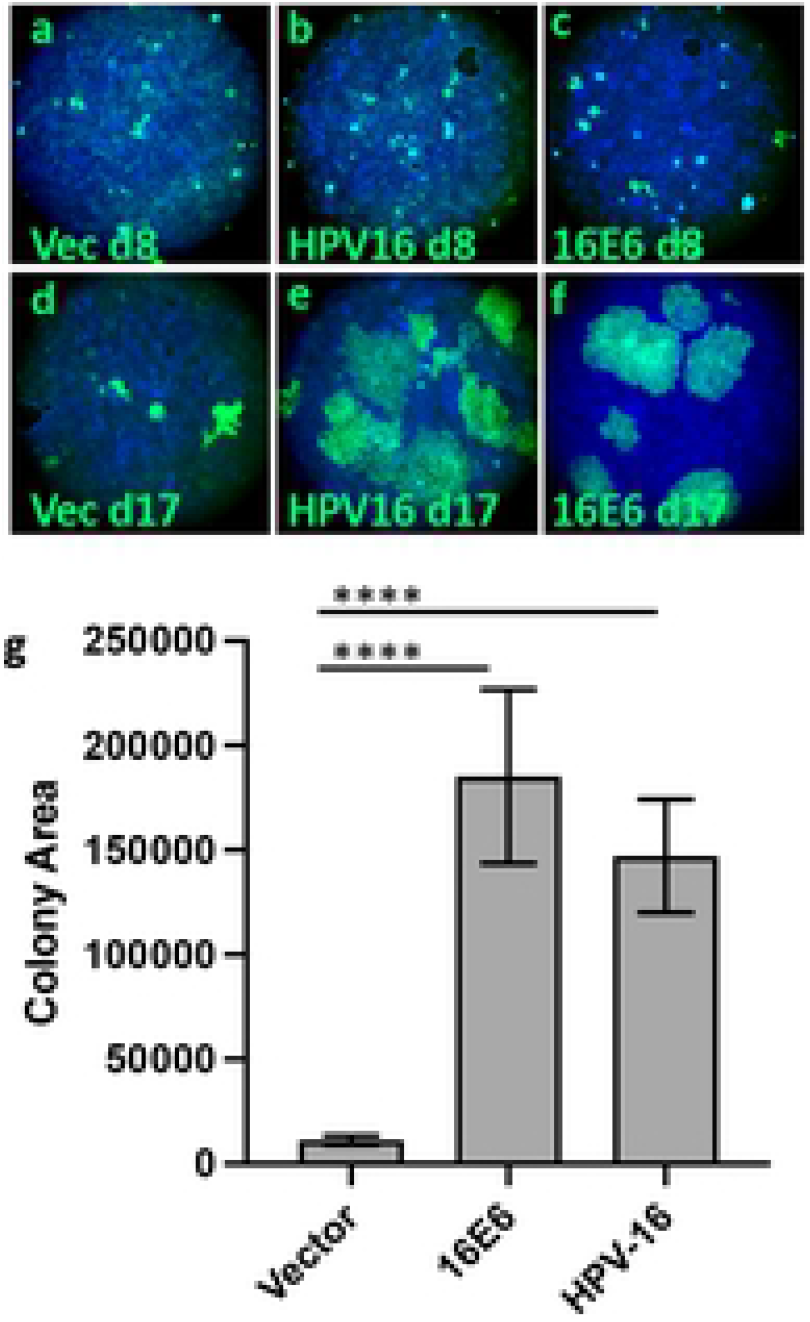
Fluorescent tagging of competing keratinocytes does not prevent cell competition by 16E6 expressing keratinocytes. EGFP-tagged NIKS cells expressing either 16E6 or the complete episomal HPV16 genome compete efficiently against un-tagged NIKS cells. Relative colony sizes on day 17 are shown in arbitrary units (g) and error bars represent standard error of the mean. **** is P<0.0001; n.s. is not significant.

To assure ourselves that simply tagging with fluorescent proteins did not impair cell competition by the surrounding keratinocytes, non-coding vector, HPV16 and 16E6 were introduced into EGFP expressing cells and set into competition against untagged keratinocytes in the same manner as shown in Fig. 2. Fig. 4 shows that EGFP labeled cells expressing either HPV16 or 16E6 competed efficiently against unlabeled keratinocytes.

We wished to ensure that the above observed results obtained in NIKS keratinocytes reflected the properties of primary keratinocytes. There is difficulty in transducing primary keratinocytes with non-transforming retroviruses as these primary cells senesce after several culture passages. To overcome this difficulty, we cultured and transduced primary keratinocytes in F media containing the rho kinase inhibitor Y-27632, which immortalizes primary keratinocytes as long as the drug is continuously present, with the cells resuming a normal cultured-cell limited lifespan when the drug is withdrawn (40). We transduced and selected primary keratinocytes in the presence of Y-27632, then continued or removed the drug after the cells were plated onto glass coverslips. Results were quite similar to results obtained in NIKS with the exception that cell spreading was larger and colonies where Y-27632 was removed were larger and less cohesive than colonies where Y-27632 was maintained throughout the assay. E6 alone enhanced colony sizes while E7 did not (Fig. 6). Western blots confirmed the expected protein expressions.

**Fig. 5.**
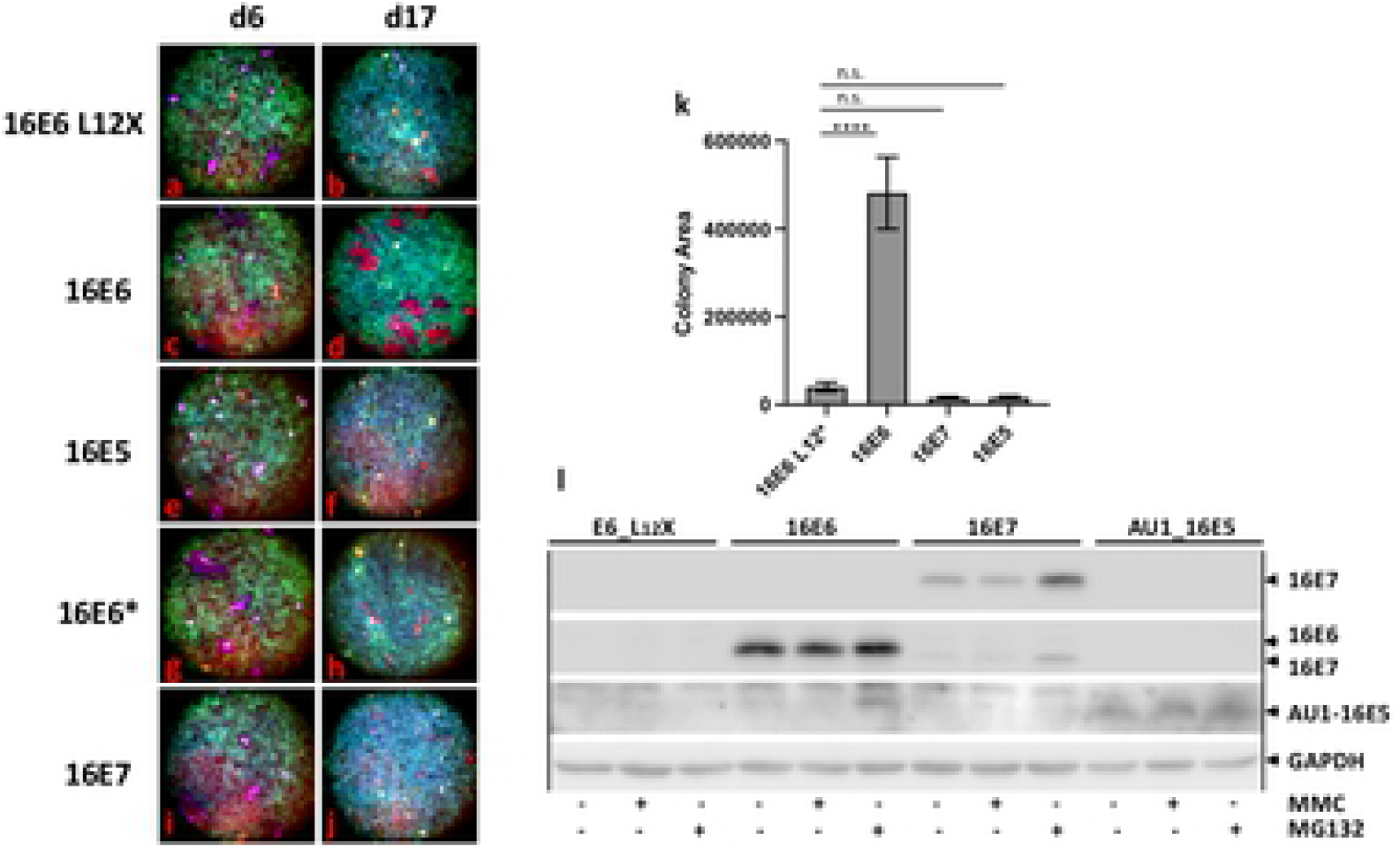
16E6 induces cell competition while 16E6*, 16E7 and 16E5 do not. The individual HPV16 oncoproteins were retrovirally transduced into Fusion-Red tagged NIKS cells and set into competition against EGFP-tagged NIKS cells as described in Fig. 1; E5 (e, f), E6 (c, d), E6* (g, h) and E7 (i, j). 16E6 with a stop codon at amino acid 12 (L12X, a and b) was the negative vector control. Plates were stained at day 6 when confluent and day 17 when super-confluent. Pictures were taken with a 4X objective and relative colony sizes ascertained at day 17 and shown in arbitrary units (k). Expression of the papillomavirus oncoproteins in the same cell lines used in this experiment is shown in part l where the blot was sequentially probed with monoclonal antibodies to E7, E6, and the AU1 epitope on E5 and finally GAPDH in that order. Day 6 colony sizes were not statistically different between the samples and are not shown. Error bars represent standard error of the mean. **** is P<0.0001; n.s. is not significant. The results shown are representative of 4 assays.

**Fig. 6.**
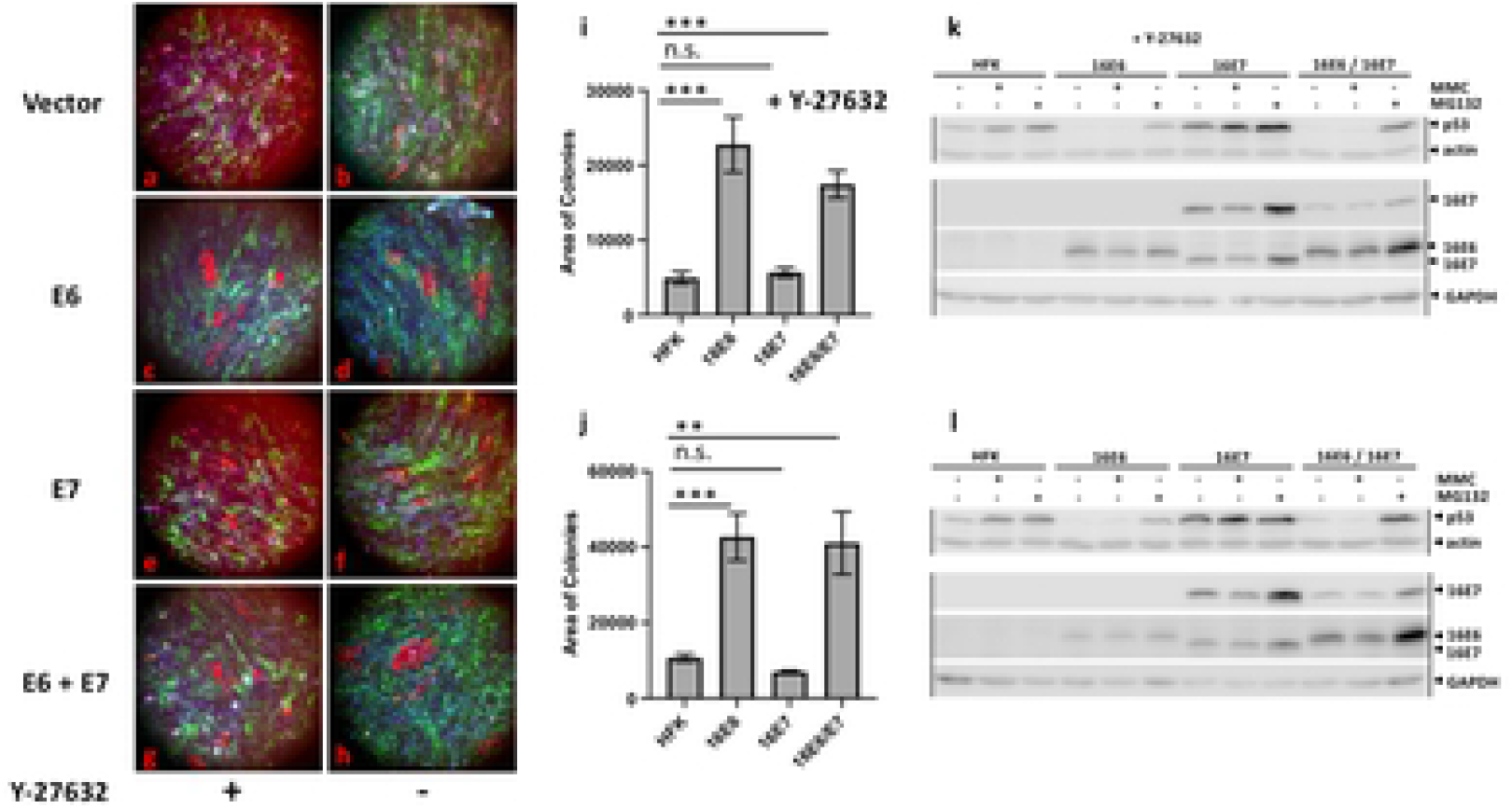
Cell competition is induced by E6 in primary keratinocytes. Primary foreskin keratinocytes maintained in the presence of the rho kinase inhibitor Y-27632 were transduced with the indicated fluorescent proteins and oncogenes as shown in Fig. 2, and seeded onto glass coverslips as described in Fig. 1. One set was maintained in media supplemented with Y-27632 (a, c, e, g, i), but in a duplicate set Y-27632 was removed after seeding onto glass coverslips (b, d, f, h, j). Cells were fixed and stained with dapi on day 21 and red colony sizes ascertained by quantitation of photomicrographs in arbitrary units. Error bars represent the variation in colony sizes, and denote standard error of the mean. *** is P<0.001; ** is P<0.0.01; n.s. is not significant. Western blots for expression of p53, actin, 16E6, 16E7, and GAPDH are shown from cells growing in the presence or absence of Y-27632 in parts k and l respectively. Results shown are representative of 4 experiments.

## Discussion

When a papillomavirus infects a basal keratinocyte, the retention of the infected keratinocyte on the basement membrane and expansion of that keratinocyte’s progeny on the basement membrane at the expense of surrounding keratinocytes is a prerequisite for the formation of a papilloma. Why papillomaviruses make papillomas at all is curious, and a priori seems unnecessary, but a papilloma would protect virus-producing cells from possibly deleterious interaction with uninfected cells. The interior of a papilloma physically segregates virus producing cells from contact with uninfected keratinocytes where that interaction might reduce the yield of virus, possibly through cell competition against cells that might be impaired by replication of virus. Our study indicates that expression of E6 in adherent cells in tissue culture confers a competitive advantage.

Key regulatory factors that influence cell competition during *Drosophila* development are proteins that in mammalian cells are found in complexes with high-risk papillomavirus E6 oncoproteins, including p53 (46–49), c-myc (50–54), and cellular PDZ proteins like SCRIB, DLG1, and dPTPMEG (the *Drosophila* homolog of PTPN3) (48, 55–59). Alteration of signaling pathways that influence cell competition in *Drosophila* are also altered by high-risk HPV E6 proteins including WNT (60, 61), pI3K/AKT and the Hippo pathway (59, 62, 63). E6 proteins have traits implicated in cell attachment and possibly cell competition; we have previously shown that the HPV16 E6 (16E6) oncoprotein enhances cell attachment and cell spreading in keratinocytes by targeted degradation of p53 (38), which could contribute to enhanced cell competition.

It is surprising that only 16E6 clearly scored in our assay and this should be cautiously interpreted. The assay we developed was designed to illuminate competition for surface area as we hypothesize would be found early in the infectious cycle. The assay is quite sensitive, and it is important to make sure that the competing cell populations are as nearly equivalent as possible with respect to drug selection markers and expressed genes; this is why we included genes with early stop codons in competing cells. Some fluorescent protein tags differed in toxicity and were unsuitable pairs for this assay. The density at which cells are plated onto the glass coverslips was important because higher initial cell density resulted in reduced adhesion of the keratinocytes, with loss of monolayer adhesion prior to the end of the assay. Different types of cell competition assays (such as those in which cells are admixed and cultured together and/or passaged together) might give rise to differing results; those assays include additional traits such as rate of cell proliferation and efficiency of cell attachment over multiple tissue culture passages. Different culture conditions such as growth factor concentrations, matrix composition, and substrate stiffness might reveal a role for either E5 or E7 that we did not observe.

The lack of clear competition conferred by E7 in our assay is surprising given that 16E7 scores much more strongly than 16E6 in both anchorage independence of murine 3T3 cells in culture, and dysplasia in transgenic mice, both being traits that so far have measured E7 traits in the absence of interaction with normal keratinocytes (26–30). It is possible that less than optimal expression levels of 16E7 from retroviruses compared to the HPV genome also might have masked a possible contribution by 16E7.

The lack of cell competition conferred by 16E5 in our assay is also in the context of isolated expression, and results might differ when expressed from the HPV16 genome together with E6. 16E5 overexpression in transgenic murine skin generates differentiation abnormalities that require EGFR signaling (64) and in transgenic mice, E5 potentiates chemical carcinogenesis (65). Again, it is noteworthy that such transgenic mouse assays (where all keratinocytes express the oncogene) excludes interaction of the oncogene-expressing cells with surrounding normal keratinocytes, and there is a natural investigator bias towards the selection of founder mice with visual abnormalities. It is possible that in cells harboring the episomal HPV16 genome that 16E5 might synergize with E6 to augment cell competition under as-yet undefined conditions, and that such a putative contribution would be lost upon integration of the genome into the cellular chromosome during progression to cancer. Neither E5 nor E7 are required for episomal maintenance of HPV16 in keratinocytes (66, 67), but E6 is required (19, 68, 69), making a genetic dissection of E6 in the context of the HPV16 genome problematic.

A genetic analysis of the effect of 16E6 on cell competition is underway, and may be complex. As noted above, 16E6 has multiple interactions with cellular proteins, many of which are candidates to mediate cell competition. 16E6 targets the degradation of p53 which has been implicated in differential competition where winner cells express lower levels of p53 (48), but while high-risk HPV E6 proteins target p53, most genera of papillomaviruses produce papillomas and do not target p53 degradation. Multiple alpha-like genera of E6 proteins do target the degradation of NHERF1 which is a negative regulator of canonical WNT signaling, and enhanced WNT signaling has been shown to enhance cell competition (70). Hippo signaling and PDZ proteins such as SCRIB and DLG that interact with 16E6 have also been implicated in modulating cell competition (57, 71), so it is possible that multiple 16E6 interactions may act together to influence competition in ways that are difficult to predict and may vary by assay conditions or the cell types infected.

There is a split in the evolutionary papillomavirus tree between virus types that encode E6 proteins that associate with the cellular E3 ubiquitin ligase E6AP (UBE3A) (72) and papillomavirus types where E6 proteins associate with MAML1 transcriptional co-activators and thereby repress Notch signaling (73–76). E6 proteins that associate with MAML1 may also associate with paxillin, a cellular adapter protein that regulates cell attachment via integrin signaling (77–80); the expression of paxillin is required for transformation by the BPV-1 E6 protein that interacts with both paxillin and MAML1 (81). Since Notch signaling controls squamous cell differentiation and carcinogenesis (82) and MAML1 is the transcriptional effector of Notch, the association of many E6 proteins with MAML1 could be a candidate interaction for influencing cell competition. Interestingly, targeted expression of a dominant negative MAML1 in the basal squamous epithelium of murine esophagus induces cell competition (83). This suggests that HPV E6 proteins that interact with MAML1 from beta and gamma genera might induce cell competition through repression of notch signaling (73, 74). If so, this would suggest the possibility that enhanced cell competition could be a broadly manifested property of E6 proteins.

Since it is now reasonable to propose that enhanced cell competition is a papillomavirus trait, a detailed genetic analysis of diverse papillomavirus oncoproteins in cell competition assays may offer broad new insights into the mechanisms by which squamous cell competition can be regulated.

